# Effects of awareness and task relevance on neurocomputational models of mismatch negativity generation

**DOI:** 10.1101/2021.02.03.429421

**Authors:** Insa Schlossmacher, Felix Lucka, Antje Peters, Maximilian Bruchmann, Thomas Straube

**Affiliations:** Institute of Medical Psychology and Systems Neuroscience, University of Münster, 48149 Münster, Germany; Otto Creutzfeldt Center for Cognitive and Behavioral Neuroscience, University of Münster, 48149 Münster, Germany; Centrum Wiskunde & Informatica, 1098XG, Amsterdam, The Netherlands; Centre for Medical Image Computing, University College London, WC1E 6BT London, United Kingdom

**Keywords:** adaptation, computational modeling, inattentional blindness, MMN, prediction

## Abstract

Detection of regularities and their violations in sensory input is key to perception. Violations are indexed by an early EEG component called the mismatch negativity (MMN) – even if participants are distracted or unaware of the stimuli. On a mechanistic level, two dominant models have been suggested to contribute to the MMN: adaptation and prediction. Whether and how context conditions, such as awareness and task relevance, modulate the mechanisms of MMN generation is unknown. We conducted an EEG study disentangling influences of task relevance and awareness on the visual MMN. Then, we estimated different computational models for the generation of single-trial amplitudes in the MMN time window. Amplitudes were best explained by a prediction error model when stimuli were task-relevant but by an adaptation model when task-irrelevant and unaware. Thus, mismatch generation does not rely on one predominant mechanism but mechanisms vary with task relevance of stimuli.

## 1. Introduction

Detecting sudden changes in our environment is of fundamental importance for perceiving and responding to altered circumstances, i.e. by adjusting actions and updating the model of the world. A typical electrophysiological phenomenon associated with deviance detection is the so-called mismatch negativity (MMN), a negative-going event-related potential (ERP) over sensory cortices in response to infrequent (deviants) compared to frequent (standards) stimuli (Näätänen et al., 1978; Stefanics et al., 2014). It has been shown that the MMN can be observed regardless whether stimuli are task-relevant or not (Alho et al., 1989; Kuldkepp et al., 2013; Näätänen et al., 1978; Schlossmacher et al., 2020) and even if participants are unaware of the stimuli of interest (Jack et al., 2017; Koelsch et al., 2006; Schlossmacher et al., 2020; Strauss et al., 2015).

While the existence of the MMN is not contested, the mechanisms behind MMN generation have been a matter of great debate (Garrido et al., 2009b; May and Tiitinen, 2010; Näätänen et al., 2005; Winkler and Czigler, 2012). Over the years, several different mechanisms of varying complexity have been put forward. One candidate is adaptation (Jääskeläinen et al., 2004; May and Tiitinen, 2010). In this framework, the observed difference between deviants and standards stems from stimulus-specific adaptation to the standards. Rare stimuli activate so-called fresh-afferents and elicit a larger response compared to the adapted standards (May and Tiitinen, 2010). Differently to the adaptation hypothesis, the memory trace hypothesis implicated that the MMN to deviants represents change detection from the build-up memory trace of the standard stimulus (Näätänen, 1992; Schröger and Winkler, 1995). Over the years the memory-trace hypothesis has been refined to also include a model adjustment aspect that relates to the updating of a model of the sensory evidence in response to unpredicted events (Winkler et al., 1996; Winkler and Czigler, 1998). Recently, model adjustment has been linked with the concept of predictive processing which has been put forward with the rise of the “Bayesian brain hypothesis” (Clark, 2013; Friston, 2005; Garrido et al., 2009b). The “Bayesian brain” describes the supposition that neural information processing relies on Bayesian principles using a generative model of the world that compares prior expectations with sensory input (Clark, 2013; Friston, 2005). From this point of view, MMN can be conceptualized as a prediction error that arises as a result of a comparison process between expected and presented stimulus. During an oddball paradigm presentation of a deviant stimulus would thus lead to a large prediction error as it violates the expectation of the more frequent standard (Stefanics et al., 2014; Winkler and Czigler, 2012). Thus, the suggested mechanisms for MMN generation differ considerably.

While in change detection schemes only the last stimulus would be relevant for the current response, an adaptation model depends on the past stimulus sequence. Similarly to adaptation, the stimulus history also plays an important role in the generation of a model of the world under a prediction account. However, while prediction is an active process that is directed in the future, adaptation is often seen as more passive (Näätänen et al., 2005). It is important to note that both adaptation and prediction can be reconciled in the predictive processing framework (Garrido et al., 2009b, 2009a; O’Shea, 2015). From this point of view, adaptation could be considered a more low-level predictive process that contributes through synaptic changes to the precision, while higher-level processes evidenced by prediction errors define the flow of information between cortical areas (Garrido et al., 2009b, 2009a). Thus from this unified perspective, observing adaptation or prediction as a mechanism in MMN generation does give important insights on the level and complexity of processing.

Computational modeling offers a unique way to investigate the underlying mechanisms by comparing predictors stemming from different models with single-trial ERP estimates (Stefanics et al., 2016). In aware conditions, computational modeling approaches underscored predictive processing as a promising mechanism during deviance processing across sensory modalities (Lieder et al., 2013; Mars et al., 2008; Ostwald et al., 2012; Stefanics et al., 2018; Weber et al., 2020). Unfortunately, until now most studies restricted their model space to two or three models excluding e.g. adaptation (Mars et al., 2008; Ostwald et al., 2012; Stefanics et al., 2018; Weber et al., 2020). However, there is evidence that under some conditions, e.g. during sleep or at low levels in the cortical hierarchy, MMN responses can at least partly be explained by adaptation (Ishishita et al., 2019; Parras et al., 2017; Strauss et al., 2015). Thus, while evidence for adaptive processes during deviance detection has been found, it is unknown whether and how experimental conditions, such as task relevance and awareness of stimuli alter the dominant mechanism at play during MMN generation.

The current study addressed these questions by investigating models underlying visual MMN under different task conditions including unawareness. One shortcoming of conventional ‘blinding’ techniques is to confound awareness of a stimulus with reporting it (Aru et al., 2012; Tsuchiya et al., 2015). In order to address this issue, participants completed a visual inattentional blindness (IB) paradigm (Pitts et al., 2012), drawing on the phenomenon that otherwise perceivable stimuli remain undetected if participants perform a distractor task and are uninformed about them (Hutchinson, 2019; Mack, 2003). This procedure allows disentangling effects of awareness and task relevance on the MMN by avoiding a trial-by-trial awareness report (Schlossmacher et al., 2020). In order to investigate how task conditions influence mechanisms of mismatch generation, we compared different computational models of single-trial amplitudes in the MMN time window during three experimental conditions ((A) unaware, (B) aware: task-irrelevant, (C) aware: task-relevant). It has been shown that MMN can be elicited during unawareness (Bekinschtein et al., 2009; Faugeras et al., 2012; Koelsch et al., 2006; Strauss et al., 2015) and that MMN as well as related components like the N1 and N2b are enhanced by attention (Auksztulewicz and Friston, 2015; Näätänen et al., 2011; Sussman et al., 2003; Sussman, 2007). Based on these findings, we expect to observe a deviance related response in all experimental conditions caused by both low-level adaptation and high-level prediction to varying degrees. We expect that mechanisms in unaware and task-irrelevant conditions rely on lower-level mechanisms like adaptation, i.e. low-level predictions, while in the task-relevant condition higher-level predictions are better suited than passive adaptation to explain deviance processing. Consequently, we propose that the relative explanatory power of predictive processing increases from unaware and task-irrelevant conditions to the task-relevant condition.

## 2. Methods

### 2.1. Participants

The sample consisted of 31 participants (9 male) aged from 18 to 35 years (*M* = 23.60, *SD* = 4.02). All had normal or corrected-to-normal vision and were right-handed. Participants volunteered and were compensated with 9 € per hour. Before starting, participants were given written instructions on the experimental task and given the opportunity to ask further questions. The study was approved by the local ethics committee and all procedures were carried out in accordance with the Helsinki declaration.

### 2.2. Apparatus

The experiment was run using MATLAB and the Psychophysics Toolbox (Brainard, 1997; Kleiner et al., 2007; Pelli, 1997). A G-Master GB2488HSU monitor at 60 Hz with a resolution of 1920 × 1080 pixels was employed for stimulus display. The viewing distance amounted to 60 cm. To respond, participants pressed the space bar and numeric keys of a standard keyboard. A chin rest was used to prevent head movements during the experiment.

### 2.3. Experimental procedure and stimulus material

Unawareness of stimuli was achieved by using an inattentional blindness paradigm (Mack, 2003; Pitts et al., 2012; Schlossmacher et al., 2020; Shafto and Pitts, 2015). In the current design, participants were presented with shapes embedded in an array of white lines presented in the background while the foreground consisted of a circling red dots that occasionally decreased in luminance. The experimental procedure included three conditions: (A) participants were either uninformed about the shapes and focused on the foreground task (unaware), or (B) were informed about the shapes but still focused on the foreground task (aware, task-irrelevant), or (C) focused on the shapes (aware, task-relevant).

Stimuli consisted of a 20 × 20 grid of white lines with a width of 0.45 degrees of visual angle (°) each, spanning 10° × 10° in total and were presented on a black background (*L_white_* = 0.35 cd/m^2^, *L_black_* = 327.43 cd/m^2^; background stimuli, see Figure 1A). Line orientation was chosen at random for each of the 400 lines comprised in the grid, i.e., a random pattern of lines was used for every presentation. Shapes were constructed by orienting lines vertically and horizontally to form a square and two rectangles centrally in the grid using 12 × 12, 8 × 16, and 16 × 8 lines, while all other lines were kept random (background stimuli, see Figure 1A). Each shape remained 100 ms on the screen followed by a random pattern presented for 700 ms, after which the next shape was presented. At all times, a red fixation cross of 0.9° × 0.9° was presented centrally. Concurrently, 12 red dots were presented on three circular paths (four on each circle) with a radius of 2.5°, 4.5°, and 6.5°, respectively (foreground stimuli; see Figure 1A for a stationary image of the dots). The dots, with radii of 0.32°, 0.41°, and 0.52°, rotated with a constant angular velocity of 1.05 radians/s. On average, every 43 s (jitter: ± 0–10 s) a randomly chosen dot slightly decreased in luminance for 500 ms (from *L* = 47.99 cd/m^2^ to *L* = 16.30 cd/m^2^; [204, 0, 0] to [114.75, 0, 0] in RGB). The rotation direction changed every 24 s on average from clockwise to counterclockwise and vice versa (jitter: ± 0–10 s). Onsets of color and rotation changes were further pseudorandomized in such a way that they never coincided with a shape onset.

**Figure 1.**
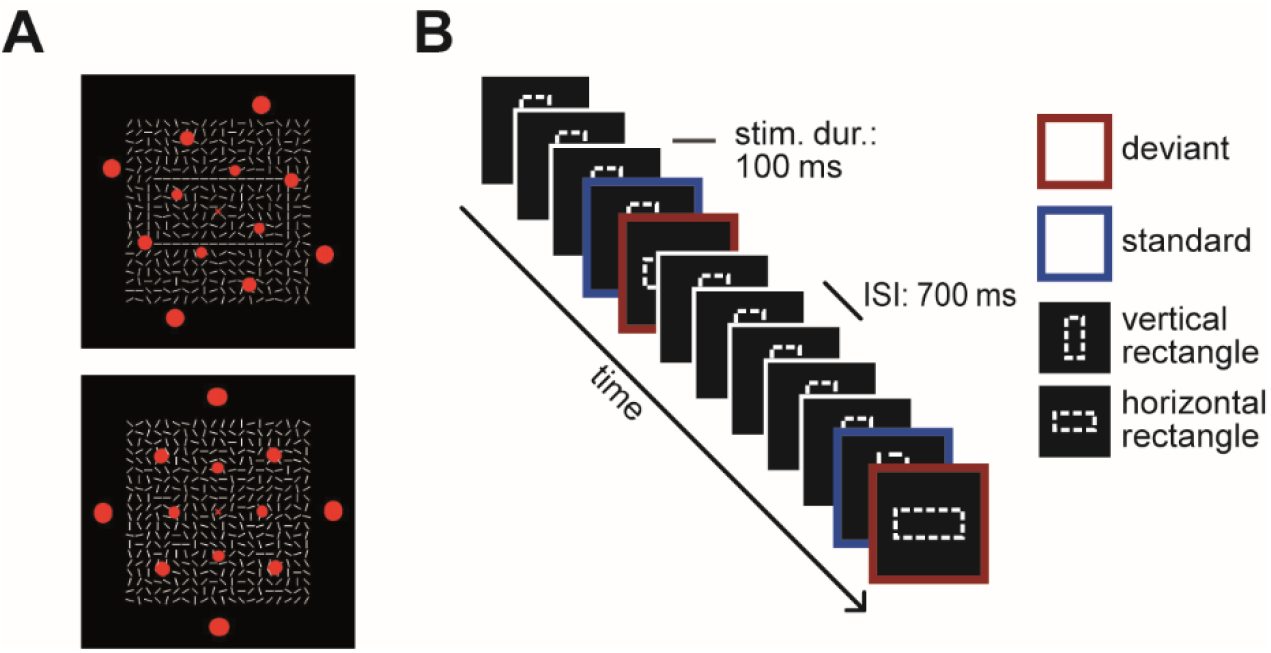
Experimental setup. (A) Example stimuli. The red dots in the foreground circled around the fixation cross and served as the task in phase A and B. In the background of the top picture a horizontal rectangle is formed out of the white line segments, in the bottom picture all lines are random. (B) Schematic of the standard oddball paradigm. Note that for the conventional analysis the standard was always the stimulus before the deviant. ISI = inter-stimulus-interval.

Shapes were presented in a standard oddball design. The standard stimulus was presented in 80% of the cases and the deviant in 20%. Stimulus presentation was further pseudorandomized that at least one standard stimulus was presented after each deviant. Horizontal and vertical rectangles served as deviant and standard and were counterbalanced across participants. On 22 randomly selected trials the shape presented was a square that served as a target in phase C. In total, 1000 stimuli were presented during each phase. The run-time of one phase amounted to 13.33 minutes (excluding breaks).

For all three experimental phases, the stimulus presentation was physically identical, while the task differed. In phase A and B, participants were instructed to press the space bar whenever they detected a luminance decrease in one of the dots. In phase C, participants’ task was to detect the squares. In phase A, participants were uninformed about the presence of the background stimuli. The difficulty of the task was designed to elicit IB in almost all of the participants. In phase B, the task was held constant, but all participants were informed about the presence of the shapes. In phase C, awareness was held constant, but participants completed a new task, which directed their attention to the shapes. Phase A was followed by either phase B or phase C counterbalanced across participants. In order to accustom participants to the tasks, phase A and phase C comprised a brief practice session in which the task difficulty was gradually increased in three steps. In phase A, the target color started with an easy-to-spot difference while only random patterns were presented until the target color used in the main experiment was reached. In phase C, the duration of shapes started with a slower presentation of 300 ms and accelerated until the shape duration of the main experiment of 100 ms was reached. For an overview of the experimental procedure, see Table 1.

**Table 1.** Overview of the experimental procedure.

| Phase | Task | Shapes task-relevant |
| --- | --- | --- |
| A | detect dot color | no |
| Awareness assessment & confidence/frequency ratings |  |  |
| B | detect dot color | no |
| Awareness assessment & confidence/frequency ratings |  |  |
| C | detect squares | yes |
| Awareness assessment & confidence/frequency ratings |  |  |

*Note:* Half of the participants completed the phases in the order ABC and the other half in the order ACB.

After completing each experimental phase, participants were given a questionnaire, asking whether or not they perceived the shapes and to describe or sketch what they saw as detailed as possible. Then, participants were asked to rate nine different shapes made of line segments (including the three shapes shown during the experiment) on how confident they were of having seen the shape (confidence rating) and how often they saw the shape (frequency rating) on a 5-point scale. The awareness questionnaire and ratings relied on Pitts and colleagues (2012) as displayed in their appendix.

### 2.4. EEG recording and preprocessing

A 128-channel BioSemi active electrode system (BioSemi B.V., Amsterdam, Netherlands) was employed to collect electrophysiological data. Electrodes were placed using the equiradial system conforming with BioSemi electrode caps. Furthermore, vertical and horizontal eye movements were recorded with two electrodes attached above and below the left eye (VEOG) and two electrodes attached to the right and left outer canthi (HEOG). Instead of ground and reference, the BioSemi EEG system uses a CMS/DRL feedback loop with two additional electrodes (for more information see: http://www.biosemi.com/faq/cms&drl.htm). Electrical potentials were recorded with a sampling rate of 512 Hz and impedances were held below 20 kΩ. A build-in analog anti-aliasing low-pass filter of 104 Hz was applied prior to digitization.

Preprocessing of the EEG data was performed using the FieldTrip toolbox (Oostenveld et al., 2010) in MATLAB. Offline filtering of the continuous data employed Butterworth filters with half-power cut-offs if not specified otherwise. Data were band-stop filtered at 49–51 Hz (roll-off: −24 dB/octave) and harmonic frequencies (up to 199–201 Hz) in order to minimize line noise. Additionally, a 59-61 Hz band-stop filter accounting for the monitor refresh rate (60 Hz) was applied. A 0.1 Hz high-pass filter (roll-off: −12 dB/octave) removed slow drifts. Then, the EEG signal was segmented into epochs of 200 ms before until 600 ms after stimulus onset. Trials containing eye blinks, muscle artifacts, and electrode jumps were manually removed based on visual inspection and bad channels were interpolated. Data were re-referenced from the CMS/DRL to a common average reference. All trials were baseline-adjusted using the average of a prestimulus interval from −200 to 0 ms.

For the oddball contrasts, trials of each subject were averaged separately for deviants and standards. We used the standard stimuli prior to deviants for the standard average allowing us to average equal amounts of stimuli per condition. This resulted in six waveforms (deviant/standard in three phases) per participant. Furthermore, deviant and standard waveforms were averaged across phases. Lastly, grand mean waveforms of the averaged data were computed.

### 2.5. Statistical analysis

To test for the vMMN, statistical analysis employed a cluster-based permutation test (Groppe et al., 2011; Maris and Oostenveld, 2007). In order to enhance its power, we chose a time interval from 150 ms to 350 ms in posterior electrodes (Stefanics et al., 2014; see Figure 2). Hypotheses were directional, leading to a one-sided cluster-based permutation test. Clusters were formed by two or more neighboring sensors (in time and space) whenever the *t*-values exceeded the cluster threshold (α = .05). The cluster mass, sum(*t*), was calculated by adding all *t*-values within a cluster. The number of permutations was set to 5000, and the significance value for testing the null hypothesis amounted to α = .05. Prior to the analysis, ERPs were down-sampled to 250 Hz and low-pass filtered at 25 Hz (roll-off: –24 dB/octave) to further enhance statistical power (Luck, 2005). In order to quantify effect sizes of the significant clusters, we averaged Cohen’s *d* for each electrode and time point. After applying the cluster-based permutation approach, cluster averages from significant clusters were computed and used in the orthogonal planned comparisons.

**Figure 2.**
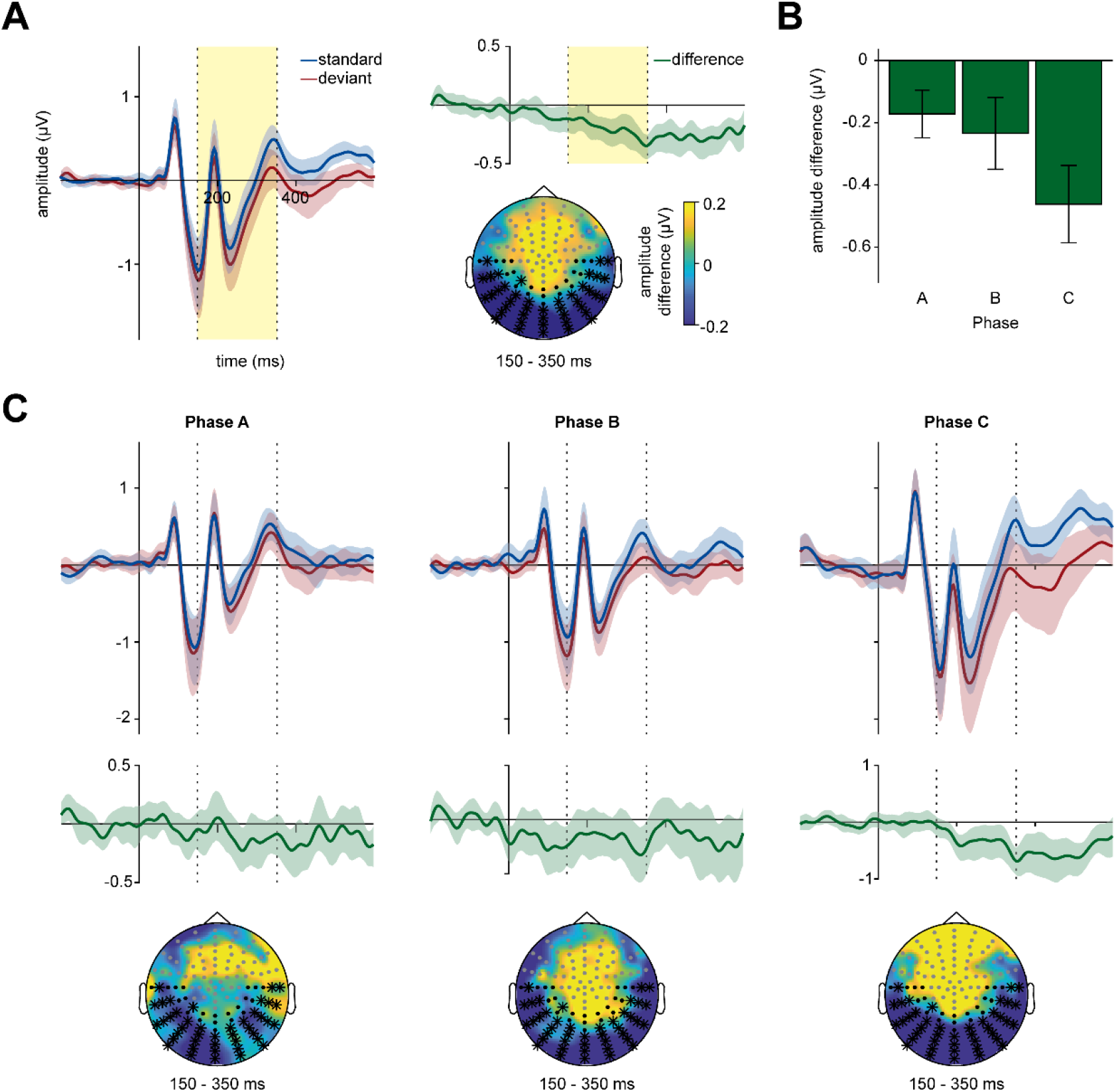
Electrophysiological measures of deviance processing. (A) VMMN effect averaged over all three phases. Black electrodes and the time interval marked by dashed lines were included in the cluster-based permutation test. Significant clusters comprised the electrodes marked as bold and the time interval marked by the light yellow box. The shaded area around ERP waveforms depicts the 95%-bootstrap confidence interval. (B) Cluster averages of vMMN in phases A, B, and C. (C) Waveforms and topographies in phases A, B and C. Error bars depict standard error of the mean.

Task performance was quantified as *d’* and reaction times of correct responses using the method for paradigms with high event rates introduced by Bendixen and Andersen (2013). This approach allows computing a false alarm rate for a continuous task with no clear distractor events. Thus, we evaluated false alarms relative to the number of non-target time intervals of the same length as the 2-seconds response interval for hits (Bendixen and Andersen, 2013). In order to probe conscious shape perception we subtracted confidence ratings for shapes included in the main experiment (hereinafter referred to as ‘shown’) from ratings for shapes not included in the main experiment (‘not shown’). In order to probe conscious perception of the oddball sequence we subtracted frequency ratings of the deviant from the standard. Rating scores, RT, *d’* and cluster averages were analyzed using repeated-measures ANOVAs and *t*-tests. Whenever sphericity was violated, the Greenhouse-Geisser correction was applied and corrected *p*-values as well as 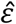-values are reported below. As some of our conclusions rely on null effects, we additionally report Bayes Factors (BF), with BF01 denoting the evidence for the null hypothesis and BF_10_ the evidence for the alternative hypothesis. Bayesian analysis relied on the R package *BayesFactor* (Morey et al., 2018), which uses a Cauchy prior scaled with 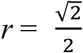 as default. We use the conventions from Jeffreys (1961) to interpret the results, that is we considered a BF > 3 as substantial evidence for either hypothesis. The cluster-based permutation was done using the FieldTrip toolbox (Oostenveld et al., 2010) in MATLAB. Other statistical tests relied on the statistics program R (R Core Team, 2015).

### 2.6. Computational modeling approach

We based our computational modeling approach on Lieder et al. (2013), who used individual single-trial estimates of the MMN and compared several different computational models like adaptation, change detection, and predictive processing accounts. However, we chose a different approach for model selection, first comparing all models with an intercept-only null model and then comparing the winning model with all other models. Furthermore, in order to account for the fitting of parameters in some of our models, we chose a 2-fold cross-validation approach with 100 repetitions (Berrar, 2019). This entailed using 50% of artifact-free single-trial vMMN estimates during the fitting procedure and using the remaining 50% to test the model fit.

Individual single-trial amplitudes were extracted using the results of the cluster-based permutation. Electrodes included in the significant vMMN cluster were averaged and an individual difference wave was computed for each participant and phase. Then, the largest negative peak of the difference wave during the time window of the significant cluster (150 – 350 ms) was determined. Last, single-trial estimates (y) were computed as the average amplitude ± 25 ms around the peak. Only artifact-free trials were used for the single-trial analysis.

We derived the single-trial trajectories of our predictors corresponding to four different alternative models, namely (1) categorical oddball, (2) change detection, (3) adaptation, and (4) precision-weighted prediction error. The categorical oddball predictor (CO) was always 1 if the stimulus was an oddball, and 0 otherwise:

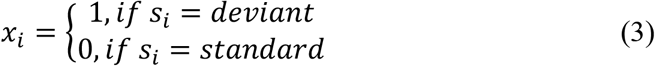

With *x_i_* representing the predictor value in trial *i*, and *s_i_* representing the stimulus type in trial *i*. We included this predictor in order to validate the findings of our classical ERP analysis as it most closely mimics the averaging procedure. The change detection predictor (CD) was 1 if the stimulus was different from the one before, and 0 otherwise, thus standards after deviants were also coded as changes:

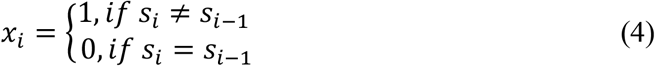

The adaptation predictor (A) was derived after Lieder et al. (2013). In this model, the response to a stimulus decays and recovers exponentially as a function of the previous stimulus sequence.

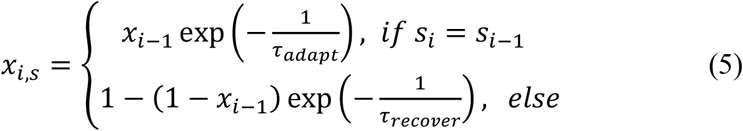

Where for each stimulus *s_i_* (here deviant and standard) a time course *x_i,s_* representing the decay and recovery of the neuronal population was computed. The predictor *x_i_* used in the final model only included the values for the stimulus shown in the experimental sequence. The additional parameters τ_adapt_ and τ_recover_ representing the rate of decay/recovery were fitted to the data using a quasi-newton optimisation algorithm implemented in *fminunc* in Matlab. We substituted values of tau that did not lie in the plausible range [0.1-200] (Lieder et al., 2013; Ulanovsky et al., 2004) with values closest to the fitted τ. We minimized the negative log-likelihood, including a parameter accounting for the observation noise similar to the HGF (see below). The precision-weighted prediction error predictor (pwPE) was derived using the freely available HGF toolbox (Mathys, 2011; Stefanics et al., 2018; Weber et al., 2020), which can be downloaded from http://www.translationalneuromodeling.org/tapas. The model builds on a process similar to Rescorla-Wagner models of reinforcement learning (Rescorla and Wagner, 1972) of the form:

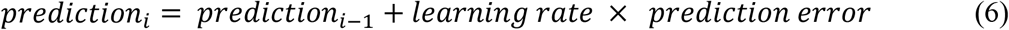

Where the prediction error is weighted by a learning rate, solving this for the pwPE gives

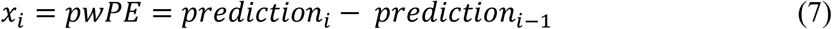

Furthermore, a hierarchical process is assumed with levels modeled as Gaussian random walks; for a description of the mathematical details of this model please refer to Mathys (2011). Here, we used the absolute second level pwPE corresponding to beliefs about the stimulus probability (Mathys, 2011) which has been shown to be relevant in mismatch processing (Stefanics et al., 2018; Weber et al., 2020). We used the HGF function *tapas_fitModel* supplied with a configuration files relying on *tapas_hgf_binary_config*, *tapas_gaussian_obs* and *tapas_quasinewton_optim_config* (TAPAS release 3.2/HGF version 5.3). In the configuration of the perceptual model, we diverged from the defaults in the following parameters. Following Stefanics et al. (2018), we fitted a two-level HGF by setting the parameter *κ_2_* to zero, thereby neglecting the environmental volatility as the oddball sequence in our experiment was highly stable. Furthermore, we changed the starting value of ω which we decreased from −3 to −6 as recommended by the HGF in order to not violate model assumptions while deriving the trajectories (*ω* = −6 (var = 16)). All other parameters were left at their respective defaults, namely 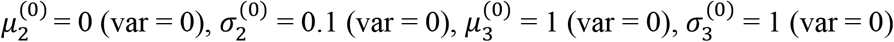, *ϑ* = −6 (var = 0), *κ*_1_ = 1. Importantly, this entailed that the initial value corresponding to the belief about the stimulus probability was set to a neutral point (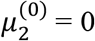, i.e. both outcomes have the same probability). For our observation model, we used *tapas_gaussian_obs* for continuous responses as a basis, but fitted the single-trial EEG estimates to the absolute second level pwPE leaving only *ω* as a free parameter. In order to accommodate the range of the single-trial estimates we set *ζ* = 4.19 corresponding to the standard deviation of single-trial amplitudes derived by using the procedure described above on the data reported in Schlossmacher et al. (2020). This procedure allowed the HGF to fit the pwPE with an appropriate decay similar to the parameters of the adaptation model.

All predictors were estimated based on the complete trial sequence (i.e., all deviants and standard stimuli); however, the parameter estimation for the adaptation and pwPE model did only take 50% of artifact-free trials into account. The remaining trials were used in the model comparison step described in the following. Using a similar approach to Mars et al. (2008), models were estimated by means of a two-level hierarchical general linear model with a random intercept and constant slope on the group level of the form

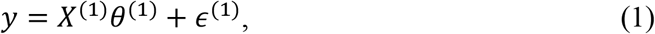

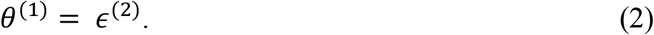

With *y* representing the concatenated single-trial estimates of all participants, *X*^(1)^ the design matrix, *θ*^(1)^ the regression parameters, and *ϵ* a random error. The design matrix *X*^(1)^ has *p* + 1 columns and *t* rows, with *p* being the number of participants and *t* being the length of the data vector *y*. 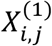 is equal to 1 if data *i* is from participant *j* and 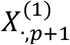 is equal to the predictor values of the tested mechanism (see below). *θ*^(1)^ is of the length *p* + 1, with the first *p* components representing the random intercepts for each participant *j* and 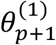 representing the regression slope. Thus, we computed one model for each phase and potential mechanism that included the data of all participants. We specifically wanted to implement a constant slope to ensure that the effects have the ‘right’ direction, i.e., a negative sign of the slope (in the current study a higher pwPE value associated with a lower single-trial amplitude consistent with the MMN).

The null model consisted only of the random intercept (i.e. 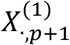 was omitted), while the alternative models additionally included predictors representing different mechanisms of mismatch generation. We computed the null model and four alternative models using the R package *lme4* (Bates et al., 2015). We compared the models using the Akaike information criterion (AIC). First, we computed Δ*AIC* = *AIC_null_* – *AIC_i_* to compare alternative models to the null model for each random test set. Then, we compared the model with the lowest AIC, i.e., the winning model, to all other models to test whether it explained the data substantially better (Δ*AIC* = *AIC_i_* – *AIC_min_*). In a last step, we averaged Δ*AIC* values to estimate model fit across all test sets and computed 95% confidence intervals. Interpretation of Δ*AIC* relied on the convention proposed by Burnham and Anderson (2004) with differences > 2 indicating substantial support for the model with the lower AIC.

### 2.7. Data and code availability statement

Data and code have been made available on the Open Science Framework accessible via https://osf.io/bjna4/.

## 3. Results

Six participants reported awareness of the shapes during phase A and were thus excluded from the analysis. With regard to the remaining 25 participants, on average, 19.41% (SD = 9.80%) of trials were excluded from the analysis due to artifacts, and on average, 0.64 (SD = 1.00) electrodes were interpolated.

### 3.1. Behavioral Data

#### 3.1.1. Task performance

Task performance quantified by *d’* did differ between phases (*F*(2,48) = 5.28, *p* = .02, 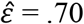, BF_10_ = 6.52) with significant better performance in phase B and C compared to phase A (all p < 0.01, all BF_10_ > 6.06), see Table 2. Comparing phase B and C, *d’* did not differ (*t*(24) = −0.92, *p* = .37, BF_01_ = 3.25). Reaction times did differ significantly between phases (*F*(2,48) = 28.42, *p* < .001, 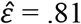, BF_10_ = 3.12×10^7^) with faster reaction times in phase C compared to both phase A and B (all *p* < .001, all BF_10_ > 7.90×10^4^), while no difference was observed between phase A and B (*t*(24) = 0.50, *p* = .62, BF_01_ = 4.24), see Table 2.

**Table 2.** Behavioral data: Performance and ratings.

| Phase | performance |  | ratings |  |
| --- | --- | --- | --- | --- |
| | $d'$ | RT | confidence | frequency |
| A | 2.30 (0.87) | 867 (141) | -0.02 (0.64) | -0.20 (0.82) |
| B | 2.72 (0.88) | 849 (120) | 2.09 (0.91) | 0.40 (0.76) |
| C | 2.94 (0.71) | 663 (63) | 2.49 (0.80) | 0.20 (0.50) |

*Note:* Reaction time (RT) measured in ms. Mean rating score differences for confidence (shown – not shown) and frequency (standard – deviant) ratings. Standard deviations are provided in parentheses.

#### 3.1.2. Confidence and frequency ratings

Repeated-measures ANOVA of rating differences (shown – not shown) indicated a significant main effect of phase for confidence (*F*(2,48) = 84.59, *p* < .001, 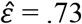, BF_10_ = 8.30×10^16^), see Table 2. Participants could significantly better differentiate shown and not shown shapes in phase C compared to phase B and in phase B and C compared to phase A (all *p* < .01, all BF_10_ > 7.76).

In the frequency rating differences (standard – deviant) a significant main effect of phase was found (*F*(2,48) = 5.42, *p* = .008, BF_10_ = 9.38), see Table 2. Participants could significantly better differentiate the frequency of standard and deviant shapes in phase C and B compared to phase A (all *p* < .05, all BF_10_ > 1.94), while no significant difference was observed between phase B and C (*t*(24) = 1.22, *p* = .23, BF_01_ = 2.43).

Importantly, *t*-tests in phase A showed no significant difference from zero for confidence (*t*(24) = −0.16, *p* = .88, BF_01_ = 4.69) indicating that ‘blind’ participants could not differentiate shown and not shown shapes. Additionally, frequency ratings did also indicate no significant difference from zero in phase A, indicating that participants were unaware of the oddball structure of the stimuli (*t*(24) = −1.22, *p* = .23, BF_01_ = 2.43).

### 3.2. EEG Data

#### 3.2.1. Electrophysiological measures of deviance processing

Averaged over all phases a significant vMMN effect was found (maximal cluster: sum(*t*) = −3081.19, *p* < .001, *d* = −0.54). This vMMN effect included the electrodes marked in the right panel of Figure 2A and lasted from 150 to 350 ms. To further substantiate that vMMN was indeed present in all phases we tested the cluster averages against zero in each phase. In phase A (*t*(24) = −2.25, *p* = .02, BF_10_ = 3.39), B (*t*(24) = −2.03, *p* = .03, BF_10_ = 2.35) and C (*t*(24) = −3.71, *p* < .001, BF_10_ = 64.81) a significant effect was observed, see Figure 2B. Furthermore, a repeated measures ANOVA with the factor phase did not indicate a significant difference in vMMN (*F*(2,48) = 1.68, *p* = .20, BF_01_ = 1.69). Testing cluster averages of the vMMN did not reveal any effects of phase order in phases B (*t*(23) = 0.58, *p* = .57, BF_01_ = 2.35) and C (*t*(23) = 0.25, *p* = .80, BF_01_ = 2.63).

#### 3.2.2. Computational modeling results

We computed the single-trial estimates in the vMMN time window around the individual peak of each participant (phase A, M = 245.31 ms, SD = 59.46 ms, min = 156.25, max = 347.66; phase B, M = 280.94 ms, SD = 61.11 ms, min = 160.16, max = 347.66; phase C, M = 262.19 ms, SD = 54.75 ms, min = 167.97, max = 347.66). Estimated averaged *θ* values for all modeled mechanisms were negative in all phases, indicating the higher the predictor the more negative the single-trial amplitude.

In phase A, the single-trial vMMN estimates were substantially better explained by the categorical oddball, precision-weighted prediction error, and adaptation model as compared to the null model (all Δ*AIC* > 2, see Figure 3B). Further, we obtained substantially more evidence for the adaptation model compared to all other models (Δ*AIC_pwPE–A_* = 6.15, 95% confidence interval [5.32 6.98], Δ*AIC_CD–A_* = 16.78 [15.75 17.79], Δ*AIC_CO–A_* = 10.06 [9.16 10.95]). In phase B, again, the categorical oddball, precision-weighted prediction error, and the adaptation model were substantially better than the null model (all Δ*AIC* > 2, see Figure 3B). Adaptation was the best model, further evidenced by substantially more evidence for this model compared to all other models (Δ*AIC_pwPE–A_* = 6.15 [5.32 6.98], Δ*AIC_CD–A_* = 16.78 [15.75 17.79], Δ*AIC_CO–A_* = 10.06 [9.16 10.95]). In phase C, all four alternative models were substantially better than the null model (all Δ*AIC* > 2, see Figure 3B). The precision-weighted prediction error was the best model, further evidenced by it being substantially better than all the other models (Δ*AIC_A–pwPE_* = 6.15 [5.32 6.98], Δ*AIC_CD–pwPE_* = 16.78 [15.75 17.79], Δ*AIC_CO–pwPE_* = 10.06 [9.16 10.95]).

**Figure 3.**
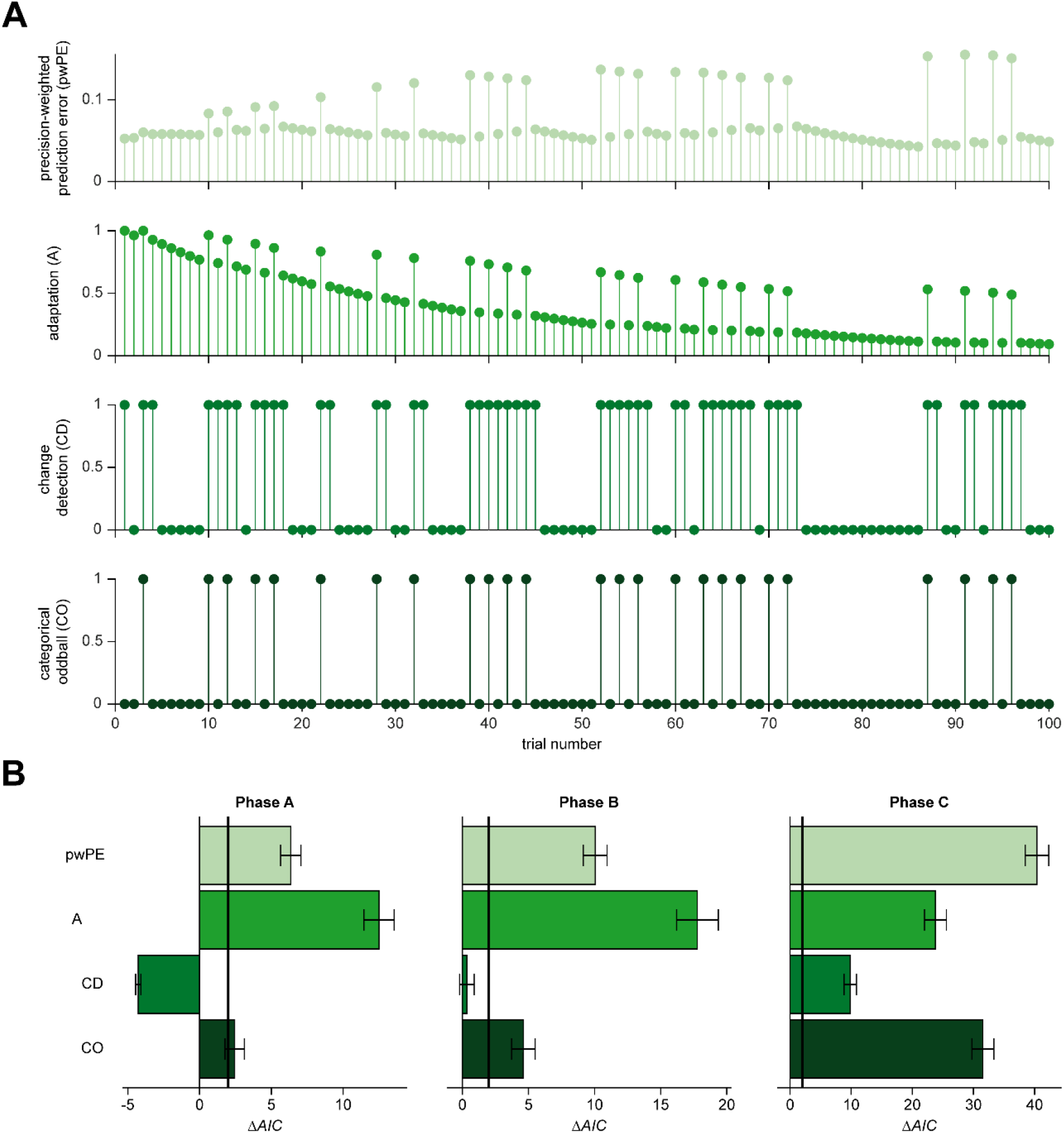
Example predictors and computational modeling results. (A) Example of single-trial predictors for one participant. The first 100 trials are depicted. (B) Alternative models compared to the null model in phase A, B and C using average Δ*AIC*. The bold line marks a Δ*AIC* of 2 which indicates substantial evidence for the alternative model after Burnham and Anderson (2004). Error bars represent 95% confidence intervals. pwPE = precision-weighted prediction error, A = adaptation, CD = change detection, CO = categorical oddball.

Fitted values of *ω*, τ_adapt_ and τ_recover_ did not differ significantly between phases (all *ps* > .05, all BF_10_ < 3) with the exception of τ_recover_ which was smaller in phase C compared to phase A (*t*(24) = 3.83, *p* < .001, BF_10_ = 41.89) and B (*t*(24) = 4.94, *p* < .001, BF_10_ = 511.33). Furthermore, we tested whether phase order influences the mechanisms by including it as a factor with an interaction term in the models of phases B and C. Models including phase order did not perform substantially better compared to their respective models without phase order effects (all Δ*AIC_without–with_* < 2).

## 4. Discussion

In the present study, we investigated the influence of awareness and task relevance on computational models of deviance processing during the visual MMN time window. Experimental manipulations had the intended effects: Participants were engaged in the tasks in all phases as evidenced by the task performance; inattentional blindness was successfully elicited in uninformed participants; we observed a vMMN in all phases. This finding agrees with the notion that the vMMN is a pre-attentive component that is elicited automatically. Interestingly, a recent study found no vMMN to low-level visual features like orientation or contrast (Male et al., 2020). In the light of this research, the vMMN observed here was probably elicited by deviance in shapes that required some form of contour integration (Hess et al., 2003) and thus more than low-level feature processing.

The predominant mechanism behind the vMMN did differ depending on experimental conditions: When stimuli were task-irrelevant, both in unaware and aware conditions, adaptation was identified as the main mechanism, while when stimuli were aware and task-relevant vMMN resembled most closely a precision-weighted prediction error. Thus, there is not only one predominant mechanism behind neuronal deviance processing but the relative contribution of mechanisms differs dependent on task settings. Thus, using MMN terminology (Kimura et al., 2009), we observed a ‘genuine’ MMN in the task-relevant condition while in the nonconscious and task-irrelevant condition deviance signals were mainly caused by refractoriness. Furthermore, we found that task relevance decreased the parameter indicating recovery from adaptation that. However, as adaptation represented only the third best model during task relevance, these results should be interpreted with caution.

In line with previous research, we found that vMMN was best explained by a prediction error (Lieder et al., 2013; Stefanics et al., 2018; Weber et al., 2020). However, this did only hold true in our task-relevant condition. Interestingly, two of these studies investigated MMN (Stefanics et al., 2018; Weber et al., 2020) during a task setting similar to our phase B, where we observed adaptation and not predictive processing. Since they did not include adaptation in their model space, a direct comparison of studies is not possible. Lieder et al. (2013) found MMN to be better explained by prediction than by adaptation in a passive auditory oddball design with a simultaneous visual task, which allows only weak control for the role of attentional focus. Another difference lies in the modality under investigation, which might have influenced results.

Our findings that unaware and distracting conditions are stronger related to adaptation as main mechanism of MMN generation fits some previous findings where mismatch processing on lower levels of the hierarchy (Ishishita et al., 2019; Parras et al., 2017) or during sleep (Strauss et al., 2015) has been shown to resemble adaptation. On the other hand, predictive processing was found on higher levels of the neuronal hierarchy and during wakefulness (Parras et al., 2017; Strauss et al., 2015). Thus, while both adaptation and prediction contribute to MMN their relative weight changes under specific contextual settings and depending on the hierarchical processing level similar to our findings. Besides, it has been shown that early vMMN responses can be explained by adaptive processes, while later portions seem to rely on memory-dependent comparison processes (Czigler et al., 2002; Kimura et al., 2009). The observed increase of predictive activity in our task-relevant phase can also be related to studies investigating effects of attention on MMN. While long considered a pre-attentive component (Näätänen et al., 2001; Sussman et al., 2014), there have been studies showing a general enhanced MMN for attended stimuli (Auksztulewicz and Friston, 2015; Sussman et al., 2003, 2014; Sussman, 2007; Woldorff et al., 1991); a finding that – while not statistically significant – can also be seen descriptively in our phase C. Furthermore, additional components like the N1 and the N2b have been found to be significantly enhanced in attended compared to unattended conditions (Näätänen et al., 2011) which might reflect increased predictive activity. This ties nicely into the observation of Auksztulewicz and Friston (2015) who found that attention is linked to enhanced top-down precision of sensory signals consistent with the predictive processing framework.

Importantly, we could show that awareness per se does not automatically elicit a dominant prediction error signal. These results are highly relevant for studies investigating MMN effects during sleep, under anesthesia, or in patients with disorders of consciousness (Bekinschtein et al., 2009; Faugeras et al., 2012; Koelsch et al., 2006; Strauss et al., 2015). In these studies, the MMN might be best explained by adaptive processes and not by predictive ones. This would fit with the observation by Strauss and colleagues (2015) who, despite observing a MMN during sleep, found that predictive processing was disrupted. Taking this up, the current results have important implications for the “Bayesian brain hypothesis”. From this point of view, the hierarchical comparison process eliciting prediction errors should be observable on all levels throughout the brain (Clark, 2013; Stefanics et al., 2014). Here, however, we found MMN to most closely resemble adaptation during unawareness and task irrelevance, while a dominant prediction error account required at least a specific amount of attentional processing of stimuli. Thus, a prediction error might not be the best model under all circumstances. Our data suggest a combination of adaptive and predictive accounts to explain the MMN shaped by attentional conditions. The more relevant the oddball stimuli the stronger is the relative contribution of predictive processes for the generation of the MMN.

As briefly sketched in the introduction, we would like to note that adaptation does not necessarily contradict the predictive brain but might be a way to accomplish it on a mechanistic level. From this point of view plastic changes in synaptic activity (i.e. adaptive processes) can be seen as a way of how the brain encodes the precision of prediction errors during predictive processing (Garrido et al., 2009a, 2009b). Adaptation can be linked to postsynaptic changes in intrinsic connections i.e. within a cortical region, while model adjustment would be mediated via extrinsic connections (Garrido et al., 2009a). Thus, adaptation might be considered a more low-level predictive process related to precision that takes place within a cortical area while higher-level processes evidenced by prediction errors allow more flexibility through extrinsic connections. This line of argument supposes that predictive processing should be observed in addition to adaptation, which was the case in the current study where both adaptation and prediction were substantially better compared to a null model in all conditions. Furthermore, it seems plausible that under conditions of unawareness and task irrelevance less extrinsic network activity and thus more adaptation is observed as has been the case in the current study.

Lastly, it is important to point out possible limitations of our study. First, inattentional blindness studies have the advantage of controlling for task relevance and awareness, but the disadvantage that only delayed reports of awareness can be used. This opens the question whether IB participants really experience blindness or rather inattentional amnesia, i.e., perceiving stimuli but swiftly forgetting them (Lamme, 2006; Wolfe, 1999). While we cannot completely rule out this possibility, one study addressing this issue found that the inability to report stimuli during IB stems from a perceptual deficit, not from memory failure (Ward and Scholl, 2015). Furthermore, the experimental setup with three consecutive phases did not allow a full counterbalancing of phase order, as the IB paradigm always has to begin with an uninformed phase to prevent conscious perception of the critical stimuli. Thus, as phase A was always the first phase, this could have potentially influenced the mechanisms observed in the following phases. However, in the light of the current findings this seems not to have been a large problem, as we observed the same mechanism in phases A and B, but different ones between phase B and C, which were counterbalanced in their order. We used a standard oddball paradigm with a rare deviant and frequent standard stimulus, which has been criticized as physical stimulus features are entangled with expectedness. While we did counterbalance the deviant across participants, a roving oddball paradigm (Baldeweg et al., 2004; Cowan et al., 1993) or appropriate control conditions (Ruhnau et al., 2012; Schröger and Wolff, 1996; Wiens et al., 2019) could alleviate this issue. The experimental tasks did manipulate attention in different ways (i.e. paying attention to distributed dots vs. paying attention to a central shape) which could potentially influenced our results. Consequently, substantiating the current findings by using different types of awareness and task manipulations as well as different oddball paradigms would be desirable.

Second, while we tried to best capture the different candidate mechanisms for MMN there might be different models that could also be included in the model space. Nevertheless, we tested four different models against a null model being able to cover the two most prominent candidates namely adaptation and prediction. Furthermore, we only modeled a monotonic relationship between predictors and EEG estimates. In addition, we used a random-intercept with a constant slope in our model comparison. While this allowed controlling the direction of the slope, not taking into account random effects on the level of the model might make the model comparison vulnerable to outliers (Stephan et al., 2009). Refined models taking into account interindividual variability and more complex coupling between EEG and predictors might be even more appropriate.

Third, we derived single-trial amplitudes individually for each participant by extracting the average amplitude around the negative peak of the difference wave. While this procedure has the advantage of specifically targeting time points relevant for mismatch processing, it might also result in different processes being covered in different participants. While individual feature selection is common in single-trial computational modeling studies (Lieder et al., 2013; Mars et al., 2008), other approaches taking all electrodes and time points into account would also be informative (Ostwald et al., 2012; Stefanics et al., 2018; Weber et al., 2020). Furthermore, we only focused on the vMMN time window and did not take into account other effects elicited during oddball paradigms, like the P3 (Polich, 2007; Verleger, 2020). The P3 has been shown to be strongly modulated by task relevance, but not by awareness (Pitts et al., 2012; Schlossmacher et al., 2020; Shafto and Pitts, 2015). As we were especially interested in how mechanisms of mismatch processing vary under different task conditions, we did not investigate the P3, which we did only expect to be elicited in our task-relevant phase (Schlossmacher et al., 2020). However, investigating in what way mechanisms of mismatch vary in this later time window depending on different task conditions including the question of whether or not the oddball is a target stimulus seems promising in future studies.

## 5. Conclusion

In summary, this EEG study investigated specific neurocomputational models of mismatch generation depending on task relevance and awareness of stimuli. A vMMN was observed in all experimental conditions. However, single-trial computational modeling showed that the adaptation model provided the best evidence in unaware and task-irrelevant conditions while a precision-weighted prediction error was the best model during task relevance. This suggests that deviance processing does not rely on either adaptation or prediction alone, but is generated by both processes whose relative contributions are dependent on task settings.

## 6. Acknowledgements

This research did not receive any specific grant from funding agencies in the public, commercial, or not-for-profit sectors. The authors declare no competing financial interests. We thank Marvin Jehn, Carolin Balloff, and Leona Rautenbach for their assistance in data collection.

